# Extracting spatial networks from capture-recapture data reveal structured habitat use within a marine mammal’s spatial range

**DOI:** 10.1101/2021.05.06.442957

**Authors:** Tyler R. Bonnell, Robert Michaud, Angélique Dupuch, Véronique Lesage, Clément Chion

## Abstract

1. Estimating the impacts of anthropogenic disturbances requires an understanding of the habitat use patterns of individuals within a population. This is especially the case when disturbances are localized within a population’s spatial range, as variation in habitat-use within a population can drastically alter the distribution of impacts.
2. Here, we illustrate the potential for multilevel multinomial models to generate spatial networks from capture-recapture data, a common data source use in wildlife studies to monitor population dynamics and habitat use. These spatial networks capture which regions of a population’s spatial distribution share similar/dissimilar individual usage patterns, and can be especially useful for detecting structured habitat use within the population’s spatial range.
3. Using simulations and 18 years of capture-recapture data from St. Lawrence Estuary (SLE) beluga, we show that this approach can successfully estimate the magnitude of similarities/dissimilarities in individual usage patterns across sectors, and identify sectors that share similar individual usage patterns that differ from other sectors, i.e., structured habitat use. In the case of SLE beluga, this method identified multiple clusters of individuals, each preferentially using restricted areas within their summer range of the SLE.
4. *Synthesis and applications.* Multilevel multinomial models can be effective at estimating spatial structure in habitat use within wildlife populations sampled by capture-recapture of individuals. Our finding of a structured habitat use within the SLE beluga summer range has direct implications for estimating individual exposures to localized stressors, such as underwater noise from shipping or other activities.

## 1. Introduction

An understanding of the spatial and temporal distribution of a species of concern is of central importance to conservation and management (Evans & Hammond, 2004). The existence of spatial structuring within populations can have important ecological and management implications. If a population as a whole can be considered as highly mixed, i.e., with individuals showing no strong patterns of home range use or sub-structuring within the wider population, then all individuals are equally likely to feel the impacts of local changes in the environment. In contrast, if the population cannot be considered to be highly mixed, and shows strong sub-structuring and site-fidelity patterns, local stressors might have a disproportionate impact on segments of the population. For example, if noise pollution increased in only one sector, in a highly mixed population all individuals would be lightly impacted, but in a spatially structured population a subset of the population would be highly impacted. These differences in spatial structuring of populations can lead to biased estimation of the likelihood and magnitude of impacts from local stressors both at the individual and population levels (DeFur et al., 2007).

Capture-recapture methods are commonly used to monitor individuals within populations, providing information on vital rates, demography, and insights into within population social mixing and habitat use (e.g., Koivuniemi et al., 2016). Photo-identification is a long-recognized method to ‘capture’ individuals with distinct markings (hereafter photo-ID data) (Urian et al., 2015), and digital photography along with high-resolution video and machine learning models to identify individuals has led to large capture-recapture datasets (Schneider et al., 2019). Novel statistical and computational methods applied to these capture-recapture datasets have enhanced the potential for quantifying within population structures through the use of social network analysis (Perryman et al., 2019; Schilds et al., 2019; Silk et al., 2021).

It is often the case, however, that efforts when collecting capture-recapture data are not evenly distributed. This is especially the case when the population under study occupies a large spatial extent, and where capture methods are not static as in the case with fixed camera traps. This variation in sampling effort can heavily bias estimates of social and spatial networks (Farine & Whitehead, 2015; Hupman et al., 2018; Whitehead, 2008). Data-stream permutations have been used to assess potential biases in network estimates from capture-recapture data when estimating networks directly from counts of individuals seen together or in the same regions (Farine, 2017; Silk et al., 2021). Alternatively, state-space models have been applied to capture-recapture data to include potential sampling biases in estimated networks when based on counts of individuals seen together (Gimenez et al., 2019). Both of these approaches build networks where individuals are the nodes, and the edges represent links between individuals. Here we propose a multilevel multinomial model approach that uses capture-recapture data to estimate spatial networks, i.e., where the nodes are spatial regions and the edges between nodes represent the magnitude of similarity in the individual using those regions. By taking this approach, it then becomes possible to quantify spatial structure in habitat use within a population’s spatial distribution.

The multilevel multinomial modeling approach that we propose to use here, does not have a large body of literature to draw on for use with capture-recapture data, but presents unique advantages (Koster & McElreath, 2017). If sampling efforts varies by region within a population’s spatial distribution, sighting probability of individuals could be greatly inflated or deflated. The use of a multilevel structure, however, allows for sighting probabilities to be nested per region and expressed in relative terms, i.e., as deviations from the mean probability of sighting. This allows the approach to identify the relative magnitude of use of a particular region for each individual. This generates a particular usage profile for each region, i.e., which individuals highly/lowly use that region (“high users” and “low users” hereafter). It is then possible to quantify how correlated the usage profiles between regions are, providing information about which regions share similar/dissimilar usage profiles. We suggest that this approach can successfully generate effort-corrected spatial networks within populations, and can help identify differential patterns in habitat use among individuals and regions.

To evaluate the performance of multilevel multinomial models at identifying spatial structuring within animal populations, we first tested the approach with simulated datasets with and without population spatial structuring. We then applied the method to observed data, using a long-term (18 years) photo-ID dataset of beluga from the St. Lawrence Estuary, Canada, and quantified spatial structuring within the population’s summer range in the St. Lawrence Estuary (SLE). Finally, we discuss how these estimates of population spatial structuring provide important information for understanding current local stressors and their potential impacts on this endangered, and declining population (Lesage, In Press).

## 2. Material and Methods

### 2.1 Data

Photo-identification boat surveys were conducted from June to October in 1989—2007 as part of an ongoing long-term study on beluga social organization. The choice of survey area on a given day was selected in a way to avoid resampling areas covered the previous days, and also according to weather conditions. When beluga were encountered, the GPS position of the research vessel was noted, and a herd follow was undertaken to photograph as many individuals as possible within the herd using a handheld camera. A herd follow was limited to three hours, with GPS location noted at least every 30 min. A detailed description of the photo-ID survey protocol is available in (Michaud, 2014). Surveys were neither systematic nor random in design, but covered various sectors of a large portion of the population’s summer distribution and a broad range of habitats. However, sampling effort was unevenly distributed across the 14 defined sectors of the summer range (Fig. 1).

**Figure 1:**
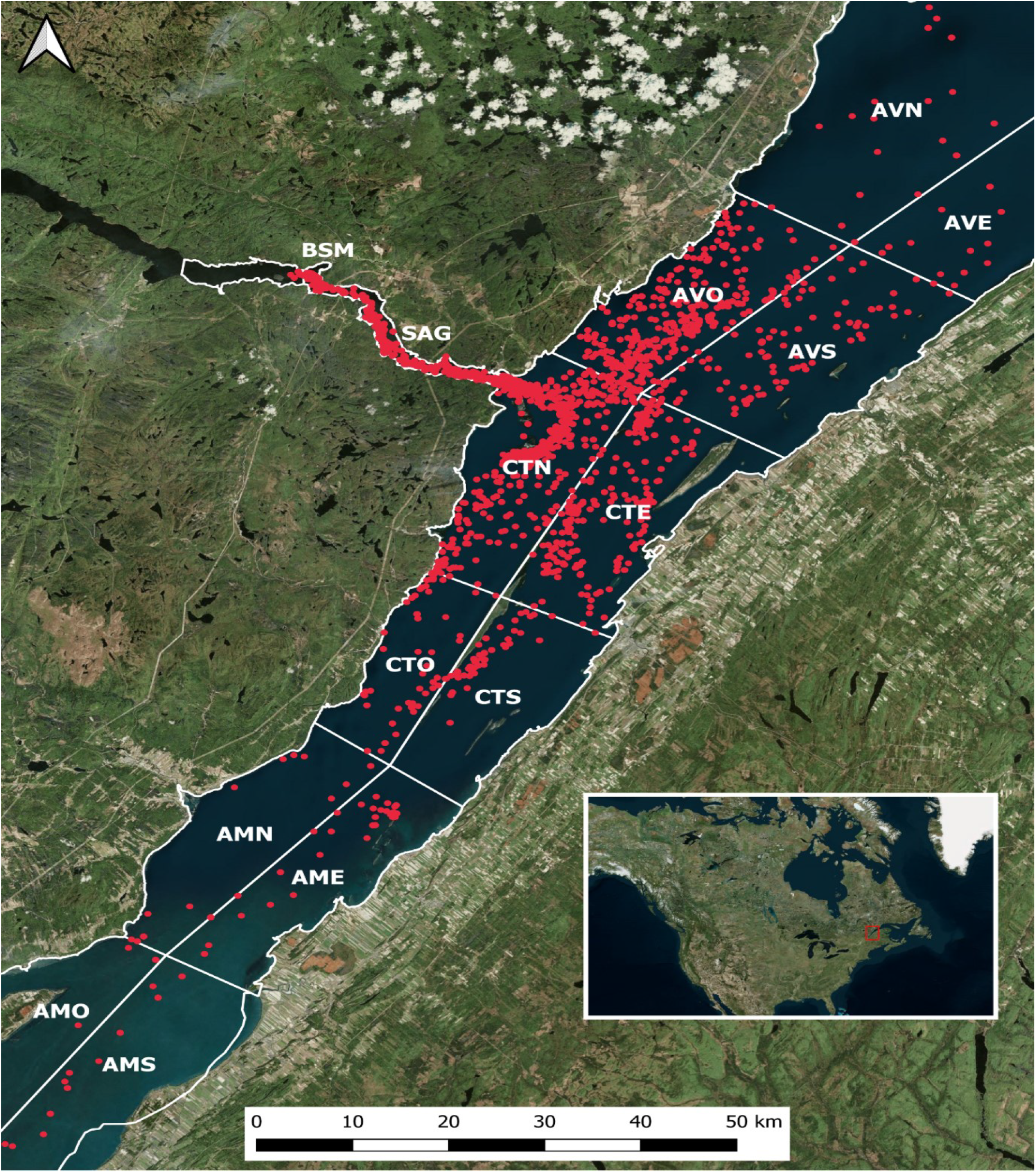
Spatial distribution of each of the 7525 photo identifications (red dots) with the 821 uniquely-identified beluga from the St. Lawrence Estuary, Canada (red square in the inset map) over our study period (1989—2007). The 14 sectors are outlined and labeled in white, and cover the summer range of the population.

Each photograph was treated using standard protocols for image selection, scoring and matching (Urian et al., 2015). Each uniquely-identified individual was attributed a resightability index ranging from 1 to 3 based on the degree of distinctiveness of markings. This resulted in a photo-identification collection of 821 unique individuals that were recaptured on average 9 times over the study period (range 1—90), and which were each associated with a GPS position and sector of initial encounter with the beluga herd. Genetically-determined sexes were available for only 29%, of the individuals included to the catalogue. From individuals with known sex, females had an estimated mean of 12 photo-IDs, and males 17 photo-IDs suggesting some bias in terms of capturability (Table S1). Given, age classes were not available for most individuals, and the low percentage of individuals with known sex, the remaining analysis focused on the population as a whole.

### 2.2 Multilevel Multinomial Model

The probability of seeing an individual in each delineated sector of the St. Lawrence Estuary was estimated using a multilevel multinomial model, where the dependent variable was the number of times an individual was captured photographically (i.e., photo-identified) in each sector. The multilevel structure of the model allowed for the estimation of both the mean probability of photo-identifying an individual in each sector, and the individual level differences in this probability by using individual ID as a random intercept. If we take, as an example, a case where the study area comprises only two sectors, then the log-odds of finding individual *i* in a sector other than the reference sector can be modeled using a multilevel multinomial following (Koster & McElreath, 2017) as:

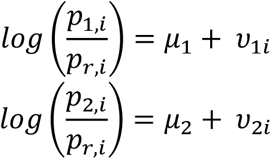

Where *p*_1,*i*_ is the probability of seeing beluga *i* in sector 1, *p*_*r,i*_ is the probability of seeing beluga *i* in the reference sector, *μ*_1_ and *μ*_2_ are the intercepts, i.e., the mean probability of seeing a beluga in sectors 1 and 2, and *v*_1*i*_ and *v*_2*i*_ are the estimated individual differences (i.e., random intercepts) from the mean probability of capture in sectors 1 and 2, respectively. The mean probabilities *μ*_1_ and *μ*_2_ represent preference/avoidance of the specified sector, while *v*_1*i*_ and *v*_2*i*_ are the sector-specific individual deviations from the mean probability of capture. As a result, it is possible to model the covariance of the individual differences between two sectors using a multivariate normal distribution:

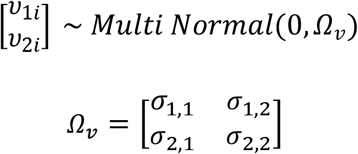

This multivariate normal distribution has a mean of 0 and a covariance matrix *Ω*_*v*_. Here the diagonal entries in the covariance matrix (*σ*_1,1_ and *σ*_2,2_) represent the magnitude of individual differences within a sector. This magnitude of individual differences identifies whether there are individual differences in the probability of being seen in a sector (i.e., high values of *σ*_1,1_ and *σ*_2,2_), or whether all individuals are equally likely to be seen (i.e., low values of *σ*_1,1_ and *σ*_2,2_). The off-diagonal entries (*σ*_2,1_ and *σ*_1,2_) are the covariance estimates between sectors, i.e., identifying sectors that share similar user profiles. By converting covariance of individual differences between sectors to correlations, this multilevel modeling approach quantifies how much information individual differences in the probability of being seen in one sector can provide about another sector. Positive correlations suggest that the high/low users in one sector are similarly high/low users in another sector, while negative correlations suggest high/low users in one sector are the low/high users in another sector.

This model can be fit with a Bayesian approach (brms package in the R environment) using a multivariate syntax: bf(y | trials(n) ∼ 1 + (1|q|ID)) + multinomial(). Here, y is a set of column vectors where each column is a sector and each row is an individual. The values in this column vector indicate how many times each individual was seen in each sector. While n is the total number of times an individual was captured photographically, q represents an arbitrary character choice that allows correlations between the estimates of random intercepts for each sector (Bürkner, 2017).

#### 2.2.1 Addition of a common reference sector

We added a preset reference category to the data (i.e., a fifteenth sector used as a reference sector) to standardize estimates of relative deviations from mean capture probability between individuals. To ensure that all individuals had the same baseline probability of being seen in the reference sector, we added the total number of captures of each individual to the new reference sector. Doing so set the probability of a capture in the reference sector to 0.5 for all individuals, i.e., equal to the probability of being captured outside of it. As the parameters in the multinomial model measured deviations away from the reference sector, we obtained standardized estimates of the relative deviations between individuals and more accurate spatial networks (Fig. S1).

#### 2.2.2 Dealing with biases in capture-recapture datasets

This multilevel multinomial model approach accounted for repeated sampling of individuals, and provided an estimate of whether some individuals were seen more or less often than the mean probability of capture in each sector (i.e. *v*_1*i*_ and *v*_2*i*_). Unlike the mean probabilities *μ*_1_ and *μ*_2_ that represent preference/avoidance of a specified sector, these estimates of individual differences from the mean probability of capture were not impacted by differential sampling among sectors. This was not the case for estimates of the mean probability of capture for each sector (i.e. *μ*_1_ and *μ*_2_), which were expected to increase in highly sampled sectors. For example, the over-sampling of the Saguenay River compared to other sectors (SAG in Fig. 1) increased the mean probability of capturing individuals in that sector. However, over-sampling of this sector was unlikely to affect the relative probability of being captured among individuals given that all individuals’ chances of being captured were likely to go up or down equally.

Similarly, potential biases due to ease of recognition, e.g., some individuals or age classes might bear more distinctive markings than others, are minimized using a multilevel multinomial approach as it focuses mainly on differences in the probability of being seen between sectors. For example, if juveniles are five times less likely to be successfully photo-identified than adults, then they might be less often represented in the photo-ID database compared to other age classes. However, the difference in distribution of these fewer photo-identified juveniles across sectors is unlikely to be impacted. For instance, if we successfully photo identified all adults 15 times and all juveniles 3 times, and if both spent twice as much of their time in the Saguenay River compared to all other sectors, then the photo-ID distribution (seen in versus outside of the Saguenay River) for adults and juveniles would be expected to be 10:5 and 2:1 respectively. In this example, the capturability varies by age class but in both age classes the probability of being captured in the Saguenay River would be twice that of the remaining sector. The adaptive partial pooling properties of multilevel models, however, leads individuals with few photo-IDs, and thus which contain less information, to be less likely to show measurable deviations from the mean probability of being captured. This means that if an age- or sex-class has very little chance of being identified by photo-ID (e.g., newborn calves or very young individuals), then they are likely to contribute less to the estimated spatial structures estimated by the multilevel multinomial approach.

By using a multilevel modeling approach, we also reduced the chance of false positives when making comparisons between many different individuals in many different sectors (i.e., problem of multiple comparisons). For example, if we were to estimate the differences in the probability of being seen in each sector separately for each individual, the risk of false positives, i.e., detecting differences where there is none, would be increased. Instead, if a multilevel approach is used to estimate the differences in probability of being seen it is possible to make effective use of partial pooling to reduce extreme values, especially in cases where the number of recaptures is not equal between individuals. Finally, by running this analysis in a Bayesian framework, we were able to place priors on the individual differences within sectors. In our case, the model was initiated assuming that there were no differences between individuals in their use of each sector, i.e., student_t(3,0,1).

### 2.3 Network analysis

Social networks are often used when visualizing and quantifying social structures within populations, with individuals often represented as nodes and their interactions as edges between these nodes (Croft et al., 2008; Farine & Whitehead, 2015). In our case, we used sectors as nodes, and the similarities in user profiles between sectors as edges (i.e., *σ*_1,2_ and *σ*_2,1_). The correlations between sectors estimated from the multilevel multinomial model can be used to create a network where the posterior predictions of each correlation parameter corresponds to an edge weight in the network. In this way, each edge has a posterior distribution and can be used to create many networks from which a distribution of network metrics can be generated, e.g., the distribution of node strength values can be calculated for each sector. The advantage of having distributions of network measures is that the measures can be readily compared, e.g., does one sector have a higher node strength than another? It is also possible to use the distribution of edge weights, and a chosen threshold (e.g., 95% credible interval), to highlight only the edges where the sign of the correlation is known with a particular range of certainty. In this paper, we used this latter approach to generate a signed network (i.e., a network with positive and negative edges), and used a simple signed-edge rule to define network communities: where a distinct community was a set of nodes that shared positive edges but no negative edges. We also made use of signed blockmodeling, an algorithm that can also be used to identify blocks of nodes, and that maximized within block positive edges and minimized within block negative edges (Doreian & Mrvar, 2015). While the signed-edge rule generally provides relatively intuitive results with simple networks, signed blockmodeling is likely to be particularly advantageous when dealing with more complex networks. The network communities detected using these two algorithms where used to identify population spatial structures.

### 2.4 Testing the multilevel multinomial modelling approach

The accuracy of the multilevel multinomial modelling approach was assessed by generating test datasets from the observed photo-ID data. We ensured that the test datasets contained the same number of unique individuals, distribution of sightings (i.e., some individuals were seen more than others), and overall number of photo-IDs as the observed dataset. We, however, varied the spatial location of individual photo-ID captures in two ways. First, to test if the proposed method correctly detected no pattern when none existed, we created a completely random test dataset by permuting the sector associated with each photo-ID in the observed dataset. The expected result was to find no correlations between sectors, given that the sectors for each photo-ID had been randomly permuted. To then test whether the proposed method could also correctly identify patterns when a known pattern existed, we generated a structured test dataset by randomly assigning each uniquely identified individual to four equally populated spatial clusters with the following and hypothetical home range of adjacent sectors: cluster 1-BSM, SAG, CTN, cluster 2- CTN, CTO, AMN, cluster 3- AVO, AVS, AVN, and cluster 4-AME, CTS, CTE. Following this, we altered the sector of where the individual photo-IDs were taken so as to fall within sectors associated with an individual’s clusters, i.e., one of their home range sectors. We did this by choosing a sector for each photo-ID based on the individual’s assigned clusters 80% of the time; a random sector was chosen for the other 20% of the time, introducing noise in the assignment of sectors. We then tested whether the model correctly identified the correlations between sectors that defined the home range of each clusters.

## 3. Results

### 3.1 Testing the multilevel multinomial modelling approach

When the multilevel multinomial model was fit to the data with sectors randomly permuted between all photo-IDs, the model found as expected no evidence for positive/negative correlations between sectors (Fig. 2a), and when we artificially created spatially distinct clusters, the model accurately estimated the correlations between sectors that defined these artificial population spatial structures (Fig. 2b). The simple signed-edge rule and blockmodeling algorithm applied to the simulated datasets both revealed the four artificially generated spatial clusters, though the blockmodeling algorithm had difficulty with the multi-membership node as it could not assign a node to two blocks (i.e., the CTN node that was shared between cluster 1 and 2).

**Figure 2:**
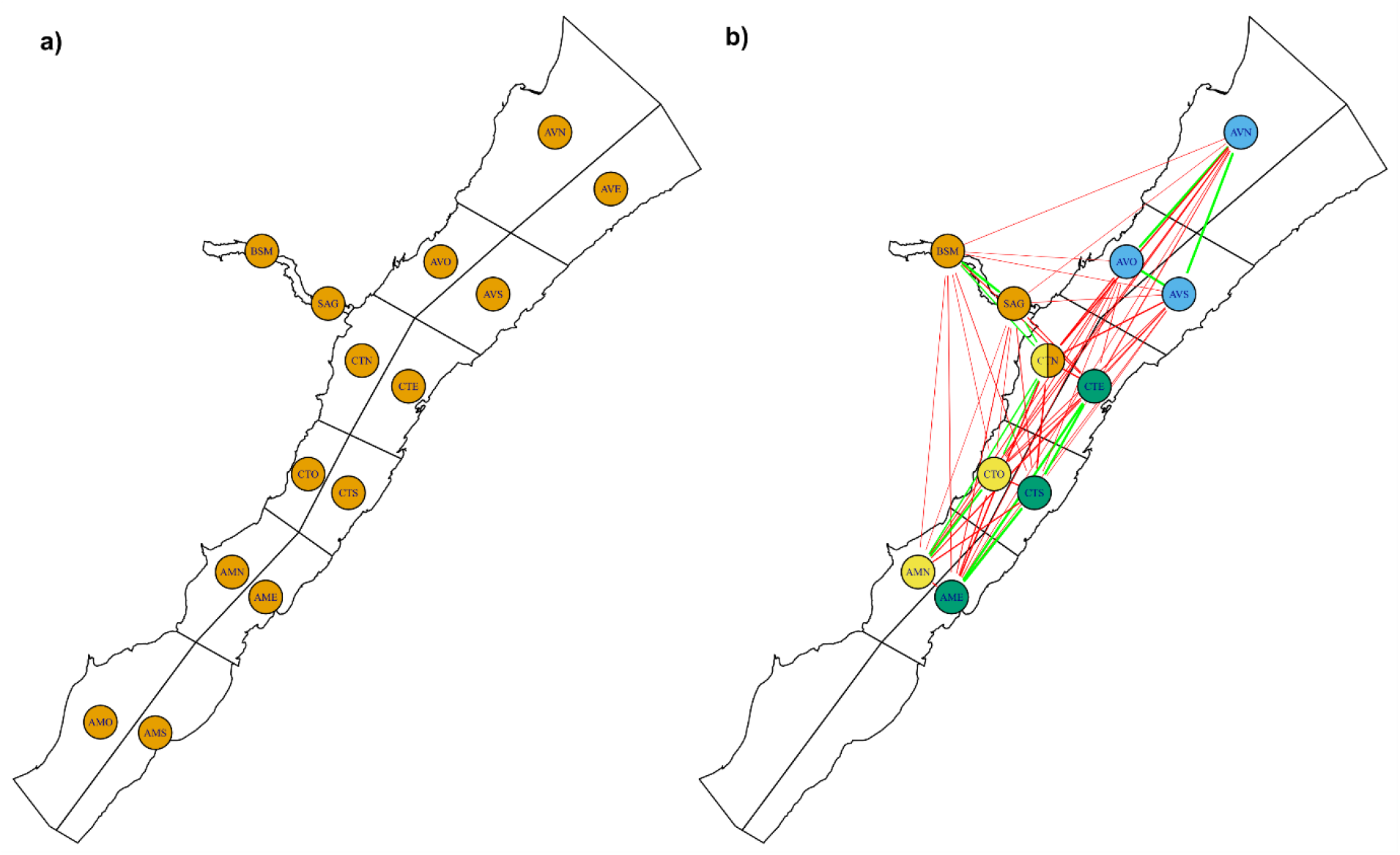
Similarity and dissimilarity between sectors in the simulated datasets: a) randomly permuted data, where there is no population spatial structure, and b) spatially structured data, where there are four distinct clusters within the population. In b) the simulated clusters are represented by color codes for each of their sectors (Note: CTN is part of the orange and yellow clusters). The green edges (lines) between two sectors signify that the sectors share high/low users, while red edges (lines) signify that they have dissimilar high/low users. The lack of an edge signifies that the high/low users of one sector does not provide information about the high/low users of other sectors.

### 3.2 Quantifying individual variation in habitat use among sectors from observed data

The multilevel multinomial model approach applied to the 18 years of observed photo-ID data indicated differences between high and low users in all sectors, though the magnitude of these individual differences varied between them (Table 1). The model also found that these individual differences were correlated between sectors (Table S2), indicating a high magnitude of similarity/dissimilarity between sectors in terms of which beluga used those sectors heavily or rarely. Taking two sectors as examples, e.g., the SAG and CTE sectors, the top 10 estimated high users of the SAG (i.e., individuals with a relatively high probability of being found there, blue dots in Fig 3a), were low users of the CTE sector (blue dots in Fig 3b).

**Table 1:**
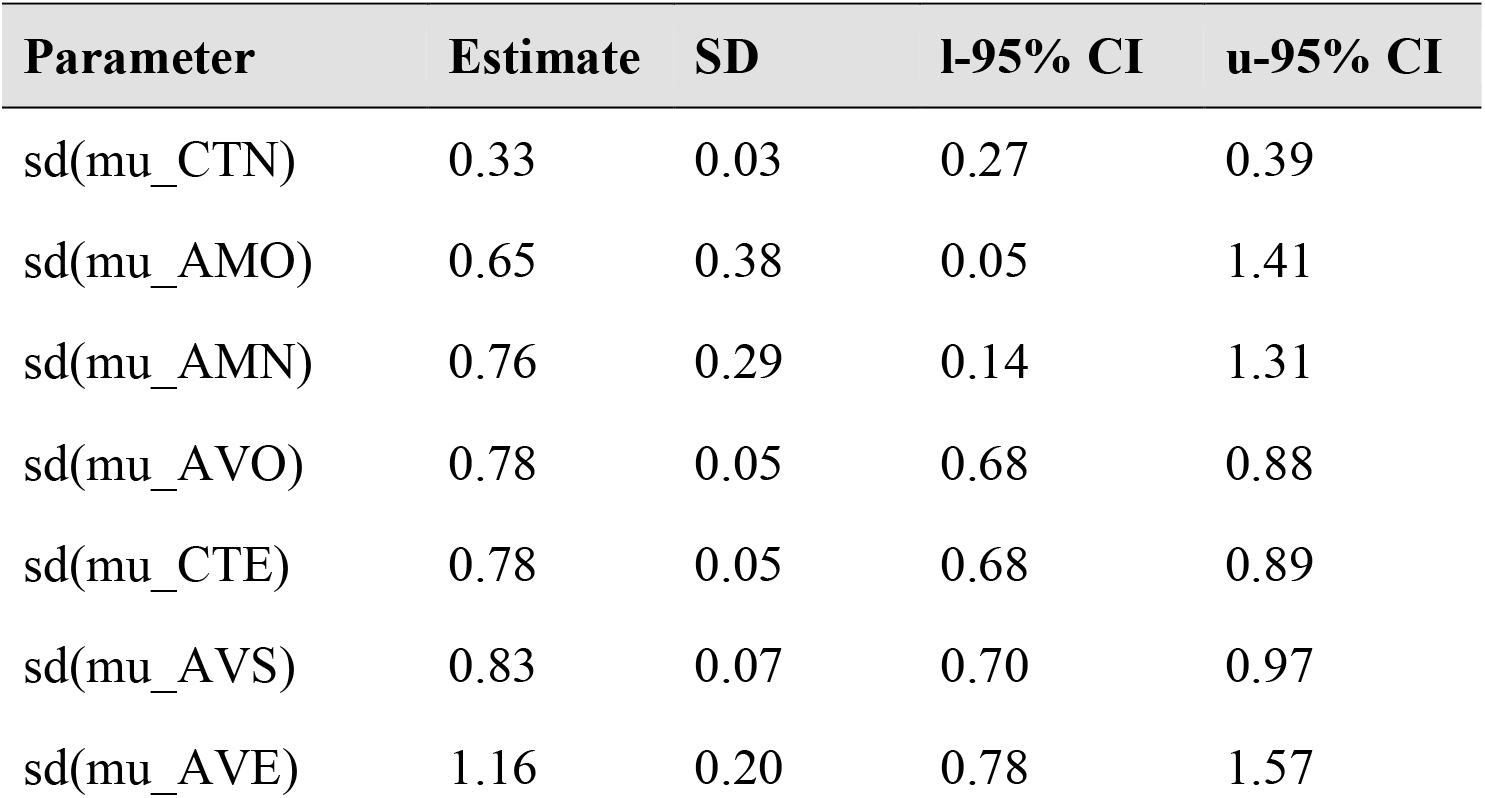
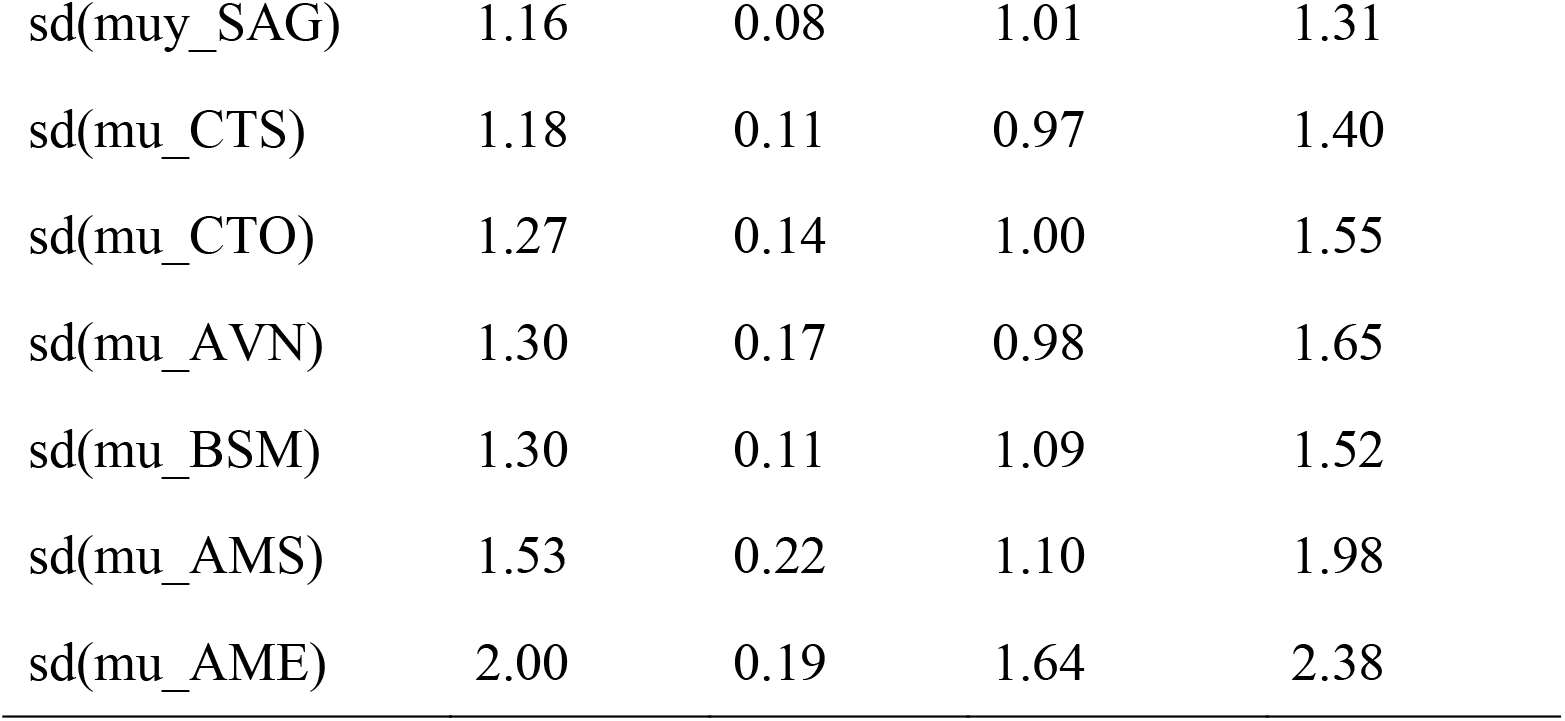
Parameter estimates from the multilevel multinomial model predicting the probability of capturing an individual by sector. Estimated magnitudes of within sector individual differences in usage (sd; e.g., *σ*_1,1_) are presented for each sector. Higher estimates indicate higher contrast between high users and low users of that sector, whereas lower estimates indicate a greater homogeneity in usage. To facilitate interpretation we have ordered the table by lowest to highest estimates of individual differences in usage, and provide the lower and upper 95% credible intervals for each estimate (e.g., l-95% CI, u-95%CI). As the number of parameters in the model is large, the overall mean by sector (i.e. *μ*_*i*_), and estimated correlations between individual differences (e.g., *σ*_2,1_) are presented in the supplementary section (Table S2).

| Parameter | Estimate | SD | l-95% CI | u-95% CI |
| --- | --- | --- | --- | --- |
| sd(mu_CTN) | 0.33 | 0.03 | 0.27 | 0.39 |
| sd(mu_AMO) | 0.65 | 0.38 | 0.05 | 1.41 |
| sd(mu_AMN) | 0.76 | 0.29 | 0.14 | 1.31 |
| sd(mu_AVO) | 0.78 | 0.05 | 0.68 | 0.88 |
| sd(mu_CTE) | 0.78 | 0.05 | 0.68 | 0.89 |
| sd(mu_AVS) | 0.83 | 0.07 | 0.70 | 0.97 |
| sd(mu_AVE) | 1.16 | 0.20 | 0.78 | 1.57 |
| sd(mu_SAG) | 1.16 | 0.08 | 1.01 | 1.31 |
| sd(mu_CTS) | 1.18 | 0.11 | 0.97 | 1.40 |
| sd(mu_CTO) | 1.27 | 0.14 | 1.00 | 1.55 |
| sd(mu_AVN) | 1.30 | 0.17 | 0.98 | 1.65 |
| sd(mu_BSM) | 1.30 | 0.11 | 1.09 | 1.52 |
| sd(mu_AMS) | 1.53 | 0.22 | 1.10 | 1.98 |
| sd(mu_AME) | 2.00 | 0.19 | 1.64 | 2.38 |

**Figure 3:**
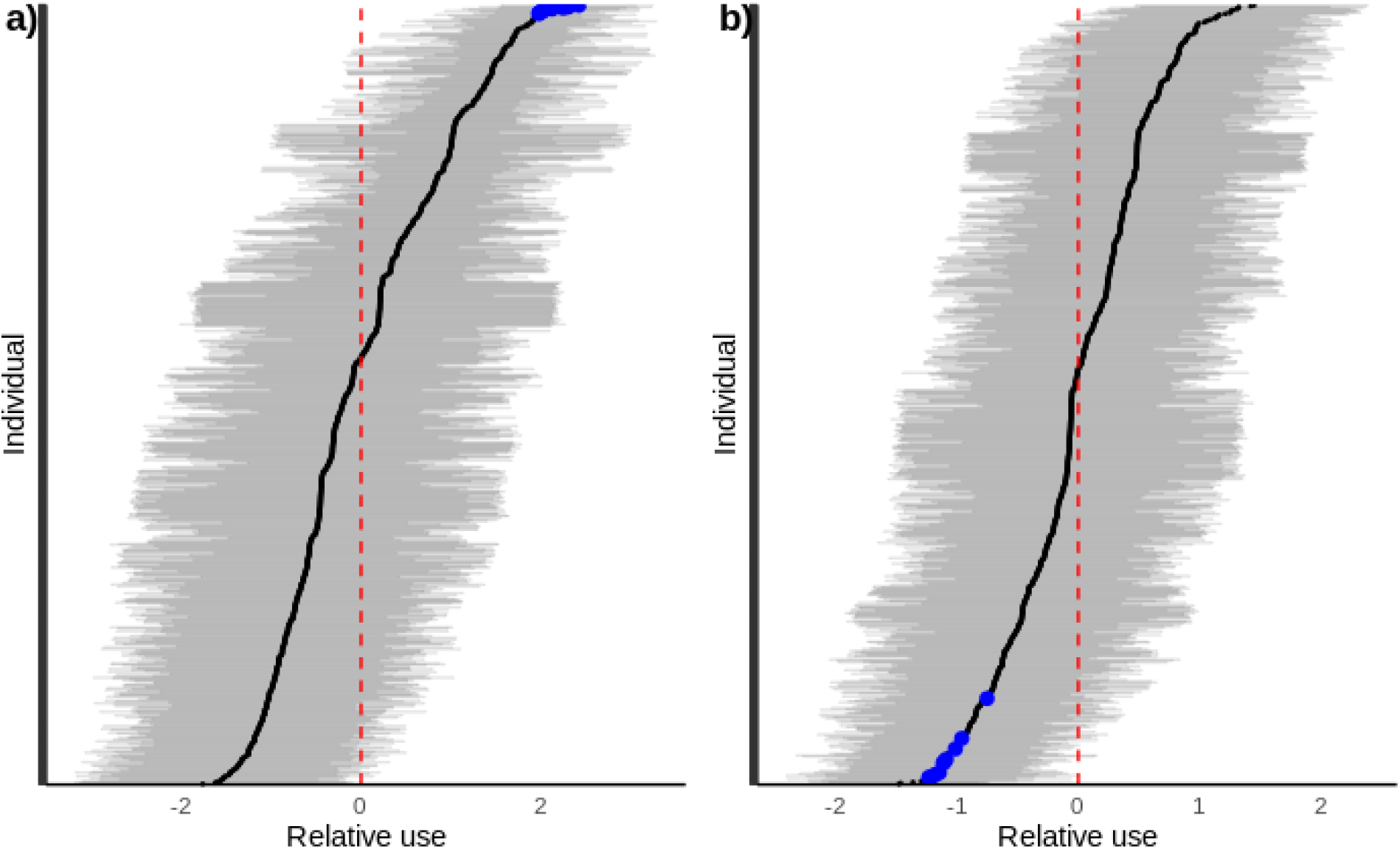
Estimate of the relative use of the SAG (a) and CTE sectors (b) by each photo-identified individual (i.e., deviation from mean use, *v*_*SAGi*_ and *v*_*CTEi*_). The values are deviations (black points) from the mean probability of recapturing individuals within a sector (red dashed line) and are on a logit scale. The horizontal grey lines represent the 95% credible interval. The estimated top 10 users of the SAG sector are represented by blue dots (panel a), and those same individuals are also highlighted in blue in the CTE sector (panel b), illustrating how correlations between sectors were estimated.

Our model indicated that the CTN sector was relatively uniformly used by all individuals (i.e., low “sd” value; Table 1) compared to other sectors. In contrast, individual differences in usage were the largest in the AME sector, with some very high/low users of that sector (Table 1).

### 3.3. Characterizing the population spatial network

Spatial patterns emerged from using the between sector correlations to generate a signed network overlaid on top of the sectors in the St. Lawrence Estuary. Applying the simple-signed rule and the blockmodeling algorithm to delineate network communities, both indicate that there are three distinct spatial clusters of individuals within the beluga summer range: the lower St. Lawrence Estuary (AVO, AVS, AVE, AVN), the Saguenay River and mouth (BSM, SAG, CTN), and the upper Estuary and eastern portion of the lower St. Lawrence Estuary (CTE, CTS, CTO, AME, AMS) (Fig. 4). In the case of AVS, however, the simple sign-rule suggested multi-membership for this sector, while the blockmodeling algorithm found AVS to be either: 1) part of the cluster containing (AVO, AVN, AVE), or 2) that the two clusters (orange and purple in fig. 4) merged into one depending on the choice of weighting parameter (i.e., emphasizing positive or negative edges).

**Figure 4:**
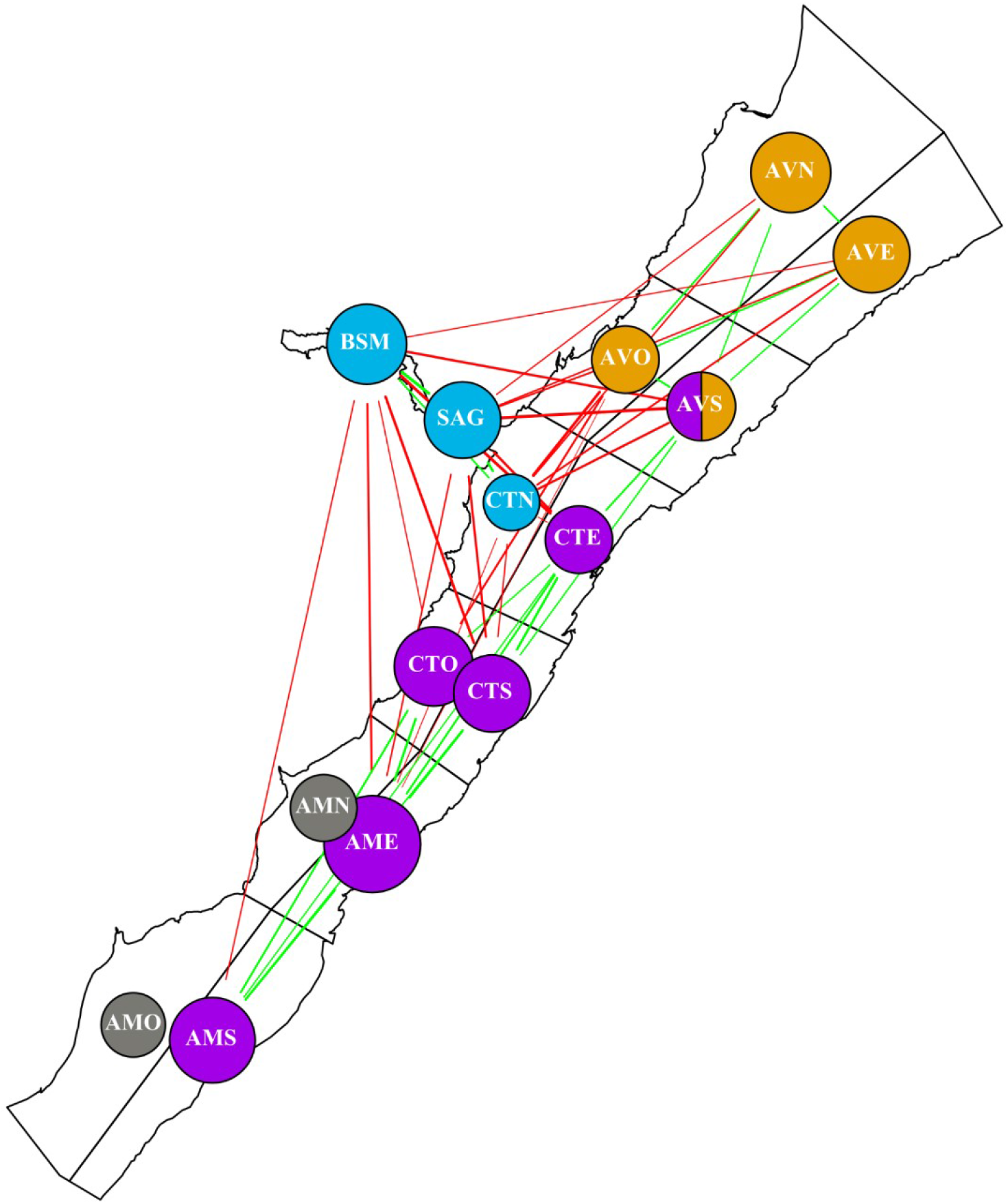
Population spatial structure characterized by similarity and dissimilarity in user profiles between sectors in the St. Lawrence Estuary beluga population. The green edges between two sectors signify that the sectors share high/low users, while red edges signify that they have dissimilar high/low users. The lack of an edge signifies that the high/low users of one sector does not provide information about the high/low users of other sectors. Nodes represent sectors, and are coloured based the cluster they belong to: i.e., shared green edges, and no shared red edges. Node sizes represent the magnitudes of individual differences in use within the sector, i.e., larger nodes suggest specialized use by a subset of the population.

## 4. Discussion

Here we’ve shown that using capture-recapture data with a multilevel multinomial modelling approach, it is possible to estimate spatial networks that can identify spatial structures within populations while controlling for sampling biases. Applying this method to data from beluga in the St. Lawrence Estuary suggests a non-random habitat use within the summer range of this population.

In particular, the use of the multilevel multinomial model provided information about within sector usage patterns of individual beluga. Our results showed that some sectors were predominantly used by a subset of individuals, while other sectors were used more uniformly by all individuals in the population. The CTN sector for instance appeared as a potential high mixing zone for the population, whereas the AME sector seemed to be used by a specific subset of the population. These findings align well with a recent study on habitat connectivity in this population exploiting a different dataset, and suggesting that the CTN sector interconnects strongly with other sectors (Ouellet et al., 2021).

The use of the multilevel multinomial model also provided information about the similarity in usage profiles across sectors. Our results across sectors add to the evidence that the beluga population cannot be assumed to be randomly mixing within its summer habitat, suggesting instead the existence of multiple spatial clusters of individuals that make use of particular sectors of the St. Lawrence Estuary and the Saguenay River. This result suggests, for example, that individuals that are repeatedly seen in the SAG sector, are also repeatedly seen in the BSM and CTN sectors, but are seen very little in the CTE and CTS sectors (Fig. 4). Given that these habitat-use patterns are constructed with all age and sex categories of beluga, and that beluga are known to segregate by age- and sex-classes during at least the summer period in the SLE and elsewhere (Loseto et al., 2006; Michaud, 1993; Smith et al., 1994), more work will be required to quantify additional population spatial structures within age/sex categories. This is crucial given that juveniles and adult females tend to show less distinctive markings compared to adult males, making captures by photo ID more difficult for this age/sex classes and reducing the amount of information they can provide when estimating population spatial networks. Ongoing photogrammetric and machine learning work to identify sex, estimate age, and facilitate individual identification will allow a more sex/age specific estimate of habitat use in this population.

Our findings about SLE beluga being spatially structured in their habitat use during summer have direct implications for estimating the impacts of anthropogenic stressors on this population. Local stressors can have a disproportionate impact on particular subsets of the larger population. Properly accounting for animal movements and population spatial structure can lead to drastically different results about impacts of individual stressors or their cumulative effects. For instance, predictions from an agent-based model of beluga and marine traffic in the STE, found that if beluga spend more time within the Saguenay River Sector they likely experience reduced exposure to noise pollution (Chion et al. Under Review). This suggests that subsets of individuals within the larger population that use this sector will have reduced noise exposure. This refuge effect, however, is predicted to be lost under scenarios where additional marine traffic is added to the Saguenay River (Chion et al. in review).

When implementing the multilevel multinomial model on other capture-recapture datasets, the use of test datasets should hold a prominent role in the analysis. The use of permutation/randomization methods to both generate spatially structured and unstructured datasets, while maintaining the sample size distribution of the original datasets, can be very valuable in helping to set model priors and to interpret the final model results. The use of permutation approaches is common in social network analysis (Croft et al., 2011; Farine, 2017), and is becoming more common in statistical workflows more generally (Gelman et al., 2013; McElreath, 2020).

## 5. Conclusions

We have introduced the use of multilevel multinomial modeling to estimate spatial networks from a capture-recapture approach that is gaining in applicability, i.e., photo-ID data. We’ve shown, using test datasets, that the proposed method is effective at detecting population spatial structures. When applied to 18 years of photo-ID data from an endangered population of beluga in the St. Lawrence Estuary, our results provide evidence that the population is composed of multiple spatial cluster of individuals with distinct habitat-use patterns. We suggest that multilevel multinomial models can be effective at extracting spatial structuring within animal populations monitored by capture-recapture sampling, contributing to impact assessments with direct implications for conservation and management.

## 6. Acknowledgment

We would like to thank all those who worked with and supported the GREMM in collecting these long term data.

## 8. Data accessibility

The permuted and simulated photo ID datasets are available on github (github.com/tbonne/photoID_multinomial), along with code used in the analysis.

## 9. Author Contributions

RM collected the data; TRB conceived the analytical methodology and performed the analysis; all authors contributed critically to the drafts and gave final approval for publication.

## References

Croft, D. P., James, R., & Krause, J. (2008). Exploring Animal Social Networks. Princeton University Press.

Croft, D. P., Madden, J. R., Franks, D. W., & James, R. (2011). Hypothesis testing in animal social networks. Trends in Ecology & Evolution, 26(10), 502–507.

DeFur, P. L., Evans, G. W., Hubal, E. A. C., Kyle, A. D., Morello-Frosch, R. A., & Williams, D. R. (2007). Vulnerability as a function of individual and group resources in cumulative risk assessment. Environmental Health Perspectives, 115(5), 817–824.

Doreian, P., & Mrvar, A. (2015). Structural balance and signed international relations. Journal of Social Structure, 16, 1.

Evans, P. G., & Hammond, P. S. (2004). Monitoring cetaceans in European waters. Mammal Review, 34(1–2), 131–156.

Farine, D. R. (2017). A guide to null models for animal social network analysis. Methods in Ecology and Evolution, 8(10), 1309–1320.

Farine, D. R., & Whitehead, H. (2015). Constructing, conducting and interpreting animal social network analysis. Journal of Animal Ecology, 84(5), 1144–1163. https://doi.org/10.1111/1365-2656.12418

Gelman, A., Carlin, J. B., Stern, H. S., Dunson, D. B., Vehtari, A., & Rubin, D. B. (2013). Bayesian data analysis. CRC press.

Gimenez, O., Mansilla, L., Klaich, M. J., Coscarella, M. A., Pedraza, S. N., & Crespo, E. A. (2019). Inferring animal social networks with imperfect detection. Ecological Modelling, 401, 69–74.

Hupman, K., Stockin, K. A., Pollock, K., Pawley, M. D. M., Dwyer, S. L., Lea, C., & Tezanos-Pinto, G. (2018). Challenges of implementing Mark-recapture studies on poorly marked gregarious delphinids. PLOS ONE, 13(7), e0198167. https://doi.org/10.1371/journal.pone.0198167

Koivuniemi, M., Auttila, M., Niemi, M., Levänen, R., & Kunnasranta, M. (2016). Photo-ID as a tool for studying and monitoring the endangered Saimaa ringed seal. Endangered Species Research, 30, 29–36.

Koster, J., & McElreath, R. (2017). Multinomial analysis of behavior: Statistical methods. Behavioral Ecology and Sociobiology, 71(9), 138.

Lesage, V. (In Press). The challenges of a small population exposed to multiple anthropogenic stressors and a changing climate: The St. Lawrence Estuary beluga. Polar Research.

Loseto, L. L., Richard, P., Stern, G. A., Orr, J., & Ferguson, S. H. (2006). Segregation of Beaufort Sea beluga whales during the open-water season. Canadian Journal of Zoology, 84(12), 1743–1751.

McElreath, R. (2020). Statistical rethinking: A Bayesian course with examples in R and Stan. CRC press.

Michaud, R. (1993). Distribution estivale du béluga du Saint-Laurent: Synthèse 1986 à 1992. Ministère des pêches et des océans, Direction de la gestion des pêches et de ….

Michaud, R. (2014). St. Lawrence Estuary beluga (Delphinapterus leucas) population parameters based on photo-identification surveys, 1989-2012. DFO Can. Sci. Advis. Sec. Res. Doc. 2013/130. iv + 27 p.

Ouellet, J.-F., Michaud, R., Moisan, M., & Lesage, V. (2021). Estimating the proportion of a beluga population using specific areas from connectivity patterns and abundance indices. Ecosphere, 00(00)(e03560). https://doi.org/10.1002/ecs2.3560

Perryman, R. J. Y., Venables, S. K., Tapilatu, R. F., Marshall, A. D., Brown, C., & Franks, D. W. (2019). Social preferences and network structure in a population of reef manta rays. Behavioral Ecology and Sociobiology, 73(8), 114. https://doi.org/10.1007/s00265-019-2720-x

Schilds, A., Mourier, J., Huveneers, C., Nazimi, L., Fox, A., & Leu, S. T. (2019). Evidence for non-random co-occurrences in a white shark aggregation. Behavioral Ecology and Sociobiology, 73(10), 138. https://doi.org/10.1007/s00265-019-2745-1

Schneider, S., Taylor, G. W., Linquist, S., & Kremer, S. C. (2019). Past, present and future approaches using computer vision for animal re-identification from camera trap data. Methods in Ecology and Evolution, 10(4), 461–470. https://doi.org/10.1111/2041-210X.13133

Silk, M. J., McDonald, R. A., Delahay, R. J., Padfield, D., & Hodgson, D. J. (2021). CMRnet: An r package to derive networks of social interactions and movement from mark– recapture data. Methods in Ecology and Evolution, 12(1), 70–75. https://doi.org/10.1111/2041-210X.13502

Smith, T. G., Hammill, M. O., & Martin, A. R. (1994). Herd composition and behaviour of white whales (Delphinapterus leucas) in two Canadian arctic estuaries. Studies of White Whales (Delphinapterus Leucas) and Narwhals (Monodon Monoceros) in Greenland and Adjacent Waters, 39, 175–184.

Urian, K., Gorgone, A., Read, A., Balmer, B., Wells, R. S., Berggren, P., Durban, J., Eguchi, T., Rayment, W., & Hammond, P. S. (2015). Recommendations for photo-identification methods used in capture-recapture models with cetaceans. Marine Mammal Science, 31(1), 298–321.

Whitehead, H. (2008). Analyzing animal societies: Quantitative methods for vertebrate social analysis. University of Chicago Press.

