## Supplementary Material for "Extracting spatial networks from capture-recapture data reveal structured habitat use within a marine mammal’s spatial range" (from another version of this preprint: version 3, 2021-06-24)

*Sex biases in captures:*

To estimate the extent to which males and females differ in their rates of capture we fit a poission regression to estimate mean rates of capture for males and females. The model results showed overdispersion and so was run with observation level random effects, which alleviated the overdispersion (Harrison, 2014). The model results suggest that males have a higher capturability (Table 1), with an estimated mean number of captures of 17 (95%CI: 15, 19) for males and 12 (95%CI: 10, 15).

Table S1: Parameter estimates from a model estimating the number of photo identification captures by sex. The reference sex category is 'unknown.'

| **Parameter** | **Estimate** | **SD** | **l-95% CI** | **u-95% CI** |
|---|---|---|---|---|
| Intercept | 1.28 | 0.04 | 1.20 | 1.35 |
| Sexe (Female) | 1.21 | 0.11 | 1.00 | 1.41 |
| Sexe (Male) | 1.56 | 0.07 | 1.42 | 1.70 |
| sd(Intercept) | 0.73 | 0.02 | 0.68 | 0.78 |

*Use of a preset reference sector:*

We use the simulated dataset with structure, presented in the main text, to fit a multilevel multinomial model with and without a preset reference sector. That is, in the model with the preset reference sector we duplicated each individual's photo-IDs and placed them in the reference sector. This results in a probability of 0.5 for being seen in the reference sector for all individuals. The results suggest that using a preset reference sector where all individuals have the same probability of being seen is required to estimate correlations between sectors and produces appropriate spatial networks (Fig S1).

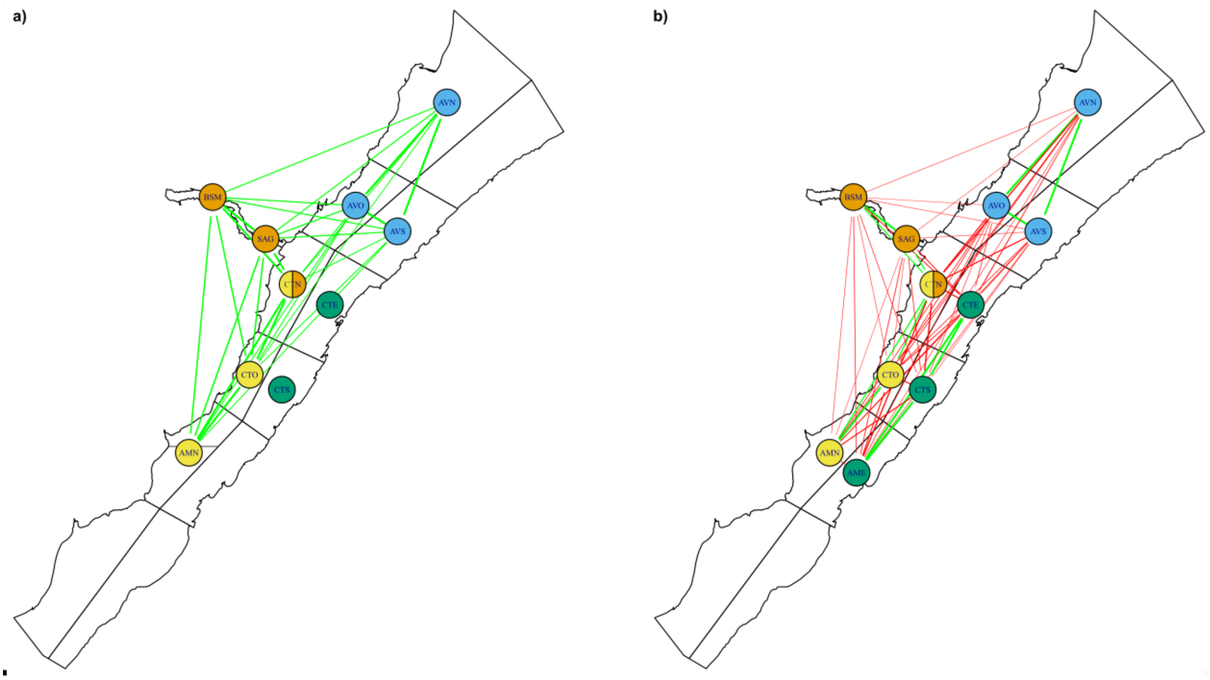


Figure S1: The estimated spatial networks for a multilevel multinormial model a) without a preset reference sector, i.e., AME is used as the reference, and b) with a preset reference sector. The green edges between two sectors signify that the sectors share high/low users, while red edges signify that they have dissimilar high/low users. The lack of an edge signifies that the high/low users of one sector does not provide information about the high/low users of other sectors. Nodes represent sectors, and are coloured based on the cluster imposed when simulating the data.

*Full model table:*

Table S2: Parameter estimates from the multilevel multinomial model predicting probability of capturing a photo-ID by sector. Estimated mean probability ($\mu_i$, logit scale), magnitude of individual differences in mean probability (e.g., $\sigma_{i,i}$), and correlation of individual differences between the different sectors (e.g., $\sigma_{i,j}$) are presented along with an estimate of their 95% credible intervals. Positive correlations suggest that the high/low users in one sector are similarly high/low users in another sector, while negative correlations suggest high/low users in one sector are the low/high users in another sector.

| Type | Parameter | Estimate | SD | l-95% CI | u-95% CI |
|---|---|---|---|---|---|
| **Mean probability $\mu_i$ (logit scale)** | | | | | |
| | mu_AME | -5.85 | 0.28 | -6.43 | -5.33 |
| | mu_AMN | -5.60 | 0.24 | -6.15 | -5.17 |
| | mu_AMO | -6.68 | 0.34 | -7.41 | -6.08 |
| | mu_AMS | -6.33 | 0.33 | -7.00 | -5.73 |
| | mu_AVE | -5.47 | 0.24 | -5.99 | -5.02 |
| | mu_AVN | -5.06 | 0.22 | -5.52 | -4.65 |
| | mu_AVO | -1.82 | 0.05 | -1.93 | -1.72 |
| | mu_AVS | -2.90 | 0.07 | -3.04 | -2.76 |
| | mu_BSM | -3.78 | 0.13 | -4.04 | -3.52 |
| | mu_CTE | -2.20 | 0.05 | -2.30 | -2.10 |
| | mu_CTN | -1.24 | 0.03 | -1.30 | -1.18 |
| | mu_CTO | -4.58 | 0.16 | -4.92 | -4.28 |
| | mu_CTS | -4.09 | 0.13 | -4.35 | -3.85 |
| | mu_SAG | -2.78 | 0.09 | -2.96 | -2.61 |
| **Individual Differences ($\sigma_{i,i}$)** | | | | | |
| | sd(mu_AME) | 2.00 | 0.19 | 1.64 | 2.38 |
| | sd(mu_AMN) | 0.76 | 0.29 | 0.14 | 1.31 |
| | sd(mu_AMO) | 0.65 | 0.38 | 0.05 | 1.41 |
| | sd(mu_AMS) | 1.53 | 0.22 | 1.10 | 1.98 |
| | sd(mu_AVE) | 1.16 | 0.20 | 0.78 | 1.57 |
| | sd(mu_AVN) | 1.30 | 0.17 | 0.98 | 1.65 |
| | sd(mu_AVO) | 0.78 | 0.05 | 0.68 | 0.88 |

| | | | | |
|---|---|---|---|---|
| sd(mu_AVS) | 0.83 | 0.07 | 0.70 | 0.97 |
| sd(mu_BSM) | 1.30 | 0.11 | 1.09 | 1.52 |
| sd(mu_CTE) | 0.78 | 0.05 | 0.68 | 0.89 |
| sd(mu_CTN) | 0.33 | 0.03 | 0.27 | 0.39 |
| sd(mu_CTO) | 1.27 | 0.14 | 1.00 | 1.55 |
| sd(mu_CTS) | 1.18 | 0.11 | 0.97 | 1.40 |
| sd(mu_SAG) | 1.16 | 0.08 | 1.01 | 1.31 |
| **Correlations between individual differences ($\sigma_{i,j}$)** | | | | |
| cor(mu_AME,mu_AMN) | 0.39 | 0.19 | -0.03 | 0.71 |
| cor(mu_AME,mu_AMO) | 0.27 | 0.24 | -0.26 | 0.67 |
| cor(mu_AMN,mu_AMO) | 0.14 | 0.24 | -0.36 | 0.57 |
| cor(mu_AME,mu_AMS) | 0.63 | 0.11 | 0.39 | 0.82 |
| cor(mu_AMN,mu_AMS) | 0.25 | 0.21 | -0.19 | 0.62 |
| cor(mu_AMO,mu_AMS) | 0.27 | 0.24 | -0.26 | 0.67 |
| cor(mu_AME,mu_AVE) | 0.18 | 0.16 | -0.14 | 0.46 |
| cor(mu_AMN,mu_AVE) | 0.15 | 0.21 | -0.28 | 0.56 |
| cor(mu_AMO,mu_AVE) | 0.10 | 0.23 | -0.36 | 0.52 |
| cor(mu_AMS,mu_AVE) | 0.05 | 0.19 | -0.32 | 0.41 |
| cor(mu_AME,mu_AVN) | -0.04 | 0.15 | -0.33 | 0.25 |
| cor(mu_AMN,mu_AVN) | -0.02 | 0.21 | -0.42 | 0.39 |
| cor(mu_AMO,mu_AVN) | 0.00 | 0.22 | -0.42 | 0.44 |
| cor(mu_AMS,mu_AVN) | -0.07 | 0.18 | -0.41 | 0.28 |
| cor(mu_AVE,mu_AVN) | 0.50 | 0.14 | 0.19 | 0.75 |
| cor(mu_AME,mu_AVO) | -0.21 | 0.10 | -0.40 | 0.00 |
| cor(mu_AMN,mu_AVO) | 0.03 | 0.19 | -0.35 | 0.40 |
| cor(mu_AMO,mu_AVO) | -0.06 | 0.21 | -0.45 | 0.36 |

| | | | | |
|---|---|---|---|---|
| cor(mu_AMS,mu_AVO) | -0.28 | 0.14 | -0.54 | 0.01 |
| cor(mu_AVE,mu_AVO) | 0.48 | 0.13 | 0.22 | 0.71 |
| cor(mu_AVN,mu_AVO) | 0.50 | 0.10 | 0.28 | 0.69 |
| cor(mu_AME,mu_AVS) | 0.13 | 0.12 | -0.10 | 0.36 |
| cor(mu_AMN,mu_AVS) | 0.21 | 0.18 | -0.16 | 0.53 |
| cor(mu_AMO,mu_AVS) | 0.13 | 0.21 | -0.31 | 0.50 |
| cor(mu_AMS,mu_AVS) | 0.03 | 0.15 | -0.26 | 0.32 |
| cor(mu_AVE,mu_AVS) | 0.40 | 0.14 | 0.12 | 0.66 |
| cor(mu_AVN,mu_AVS) | 0.38 | 0.12 | 0.14 | 0.60 |
| cor(mu_AVO,mu_AVS) | 0.54 | 0.08 | 0.38 | 0.69 |
| cor(mu_AME,mu_BSM) | -0.57 | 0.09 | -0.75 | -0.38 |
| cor(mu_AMN,mu_BSM) | -0.37 | 0.17 | -0.66 | 0.00 |
| cor(mu_AMO,mu_BSM) | -0.22 | 0.23 | -0.62 | 0.26 |
| cor(mu_AMS,mu_BSM) | -0.37 | 0.13 | -0.62 | -0.10 |
| cor(mu_AVE,mu_BSM) | -0.34 | 0.13 | -0.59 | -0.06 |
| cor(mu_AVN,mu_BSM) | -0.22 | 0.12 | -0.44 | 0.02 |
| cor(mu_AVO,mu_BSM) | -0.16 | 0.08 | -0.32 | 0.01 |
| cor(mu_AVS,mu_BSM) | -0.66 | 0.07 | -0.80 | -0.51 |
| cor(mu_AME,mu_CTE) | 0.54 | 0.10 | 0.34 | 0.72 |
| cor(mu_AMN,mu_CTE) | 0.34 | 0.18 | -0.04 | 0.65 |
| cor(mu_AMO,mu_CTE) | 0.21 | 0.23 | -0.29 | 0.60 |
| cor(mu_AMS,mu_CTE) | 0.38 | 0.14 | 0.10 | 0.62 |
| cor(mu_AVE,mu_CTE) | 0.07 | 0.14 | -0.21 | 0.36 |
| cor(mu_AVN,mu_CTE) | -0.11 | 0.12 | -0.35 | 0.14 |
| cor(mu_AVO,mu_CTE) | -0.10 | 0.08 | -0.26 | 0.07 |
| cor(mu_AVS,mu_CTE) | 0.52 | 0.08 | 0.34 | 0.67 |

| | | | | |
|---|---|---|---|---|
| cor(mu_BSM,mu_CTE) | -0.80 | 0.06 | -0.89 | -0.68 |
| cor(mu_AME,mu_CTN) | -0.26 | 0.12 | -0.49 | -0.03 |
| cor(mu_AMN,mu_CTN) | -0.24 | 0.19 | -0.59 | 0.16 |
| cor(mu_AMO,mu_CTN) | -0.12 | 0.21 | -0.50 | 0.32 |
| cor(mu_AMS,mu_CTN) | -0.09 | 0.15 | -0.39 | 0.21 |
| cor(mu_AVE,mu_CTN) | -0.53 | 0.13 | -0.77 | -0.26 |
| cor(mu_AVN,mu_CTN) | -0.44 | 0.12 | -0.67 | -0.18 |
| cor(mu_AVO,mu_CTN) | -0.75 | 0.07 | -0.86 | -0.60 |
| cor(mu_AVS,mu_CTN) | -0.63 | 0.09 | -0.79 | -0.44 |
| cor(mu_BSM,mu_CTN) | 0.47 | 0.09 | 0.28 | 0.64 |
| cor(mu_CTE,mu_CTN) | -0.25 | 0.10 | -0.46 | -0.05 |
| cor(mu_AME,mu_CTO) | 0.63 | 0.09 | 0.43 | 0.79 |
| cor(mu_AMN,mu_CTO) | 0.28 | 0.20 | -0.15 | 0.63 |
| cor(mu_AMO,mu_CTO) | 0.16 | 0.22 | -0.30 | 0.56 |
| cor(mu_AMS,mu_CTO) | 0.55 | 0.13 | 0.27 | 0.78 |
| cor(mu_AVE,mu_CTO) | -0.12 | 0.17 | -0.45 | 0.22 |
| cor(mu_AVN,mu_CTO) | -0.24 | 0.15 | -0.54 | 0.07 |
| cor(mu_AVO,mu_CTO) | -0.55 | 0.09 | -0.72 | -0.36 |
| cor(mu_AVS,mu_CTO) | -0.16 | 0.12 | -0.39 | 0.07 |
| cor(mu_BSM,mu_CTO) | -0.33 | 0.11 | -0.53 | -0.12 |
| cor(mu_CTE,mu_CTO) | 0.39 | 0.11 | 0.17 | 0.60 |
| cor(mu_CTN,mu_CTO) | 0.19 | 0.12 | -0.06 | 0.42 |
| cor(mu_AME,mu_CTS) | 0.70 | 0.08 | 0.53 | 0.85 |
| cor(mu_AMN,mu_CTS) | 0.42 | 0.19 | 0.00 | 0.73 |
| cor(mu_AMO,mu_CTS) | 0.21 | 0.23 | -0.27 | 0.61 |
| cor(mu_AMS,mu_CTS) | 0.42 | 0.14 | 0.14 | 0.67 |

| | | | | |
|---|---|---|---|---|
| cor(mu_AVE,mu_CTS) | 0.16 | 0.16 | -0.15 | 0.45 |
| cor(mu_AVN,mu_CTS) | 0.06 | 0.14 | -0.23 | 0.33 |
| cor(mu_AVO,mu_CTS) | -0.04 | 0.10 | -0.23 | 0.15 |
| cor(mu_AVS,mu_CTS) | 0.42 | 0.11 | 0.20 | 0.61 |
| cor(mu_BSM,mu_CTS) | -0.72 | 0.08 | -0.85 | -0.55 |
| cor(mu_CTE,mu_CTS) | 0.64 | 0.09 | 0.46 | 0.80 |
| cor(mu_CTN,mu_CTS) | -0.37 | 0.11 | -0.57 | -0.14 |
| cor(mu_CTO,mu_CTS) | 0.48 | 0.11 | 0.25 | 0.68 |
| cor(mu_AME,mu_SAG) | -0.43 | 0.09 | -0.61 | -0.24 |
| cor(mu_AMN,mu_SAG) | -0.36 | 0.17 | -0.66 | 0.00 |
| cor(mu_AMO,mu_SAG) | -0.17 | 0.22 | -0.56 | 0.29 |
| cor(mu_AMS,mu_SAG) | -0.22 | 0.13 | -0.47 | 0.03 |
| cor(mu_AVE,mu_SAG) | -0.44 | 0.12 | -0.68 | -0.19 |
| cor(mu_AVN,mu_SAG) | -0.37 | 0.10 | -0.57 | -0.16 |
| cor(mu_AVO,mu_SAG) | -0.48 | 0.06 | -0.60 | -0.35 |
| cor(mu_AVS,mu_SAG) | -0.80 | 0.06 | -0.90 | -0.68 |
| cor(mu_BSM,mu_SAG) | 0.83 | 0.04 | 0.74 | 0.91 |
| cor(mu_CTE,mu_SAG) | -0.68 | 0.06 | -0.79 | -0.55 |
| cor(mu_CTN,mu_SAG) | 0.67 | 0.07 | 0.52 | 0.80 |
| cor(mu_CTO,mu_SAG) | -0.12 | 0.11 | -0.32 | 0.09 |
| cor(mu_CTS,mu_SAG) | -0.62 | 0.08 | -0.77 | -0.45 |

*Note on interpreting low sd within sectors:*

Similar to the CTN sector, the sector AMO is also estimated to have a low magnitude of individual differences. However, it has a large uncertainty in this estimate. This highlights that it is possible to have little individual differences in a sector due to either: 1) limited data, resulting in all individuals being pooled to the mean value, and 2) limited data is not a factor, but individuals are using this sector relatively equally. Care should therefore be taken when interpreting the magnitude of individual differences, nevertheless, the estimated uncertainty around magnitude estimates is one way to help identify sectors with limited data.

*References:*

Harrison, X. A. (2014). Using observation-level random effects to model overdispersion in count data in ecology and evolution. *PeerJ*, *2*, e616.
